# Slope of the power spectral density flattens at low frequencies (<150 Hz) with healthy aging but also steepens at higher frequency (>200 Hz) in human electroencephalogram

**DOI:** 10.1101/2023.02.15.528644

**Authors:** Srishty Aggarwal, Supratim Ray

## Abstract

Brain signals such as electroencephalogram (EEG) often show oscillations at various frequencies, which are represented as distinct “bumps” in the power spectral density (PSD) of these signals. In addition, the PSD also shows a distinct reduction in power with increasing frequency, which pertains to aperiodic activity and is often termed as the “1/f” component. While a change in periodic activity in brain signals with healthy aging and mental disorders has been reported, recent studies have shown a reduction in the slope of the aperiodic activity with these factors as well. However, these studies only analysed PSD slopes over a limited frequency range (<100 Hz). To test whether the PSD slope is affected over a wider frequency range with aging and mental disorder, we collected EEG data with high sampling rate (2500 Hz) from a large population of elderly subjects (>49 years) who were healthy (N=217) or had mild cognitive impairment (MCI; N=11) or Alzheimer’s Disease (AD; N=5), and analysed the PSD slope till 800 Hz. Consistent with previous studies, the 1/f slope up to ~150 Hz reduced with healthy aging. Surprisingly, we found the opposite at higher frequencies (>200 Hz): the slope increased with age. This result was observed in all electrodes, for both eyes open and eyes closed conditions, and for different reference schemes. Slopes were not significantly different in MCI/AD subjects compared to age and gender matched healthy controls. Overall, our results constrain the biophysical mechanisms that are reflected in the PSD slopes in healthy and pathological aging.

**Significance Statement:** Aperiodic activity in the brain is characterized by measuring the slope of the power spectrum of brain signals. This slope has been shown to flatten with healthy aging, suggesting an increase in some sort of “neural noise”. However, this flattening has been observed only over a limited frequency range (<150 Hz). We found that at higher frequencies (>200 Hz), the opposite happens: the slope steepens with age. This occurs at all electrodes, irrespective of state and referencing techniques. However, the slope is unchanged in subjects with early Alzheimer’s Disease (AD) and their controls. Our results shed new light on the properties of neural noise and the neurophysiological processes affecting AD and the aperiodic activity.

## Introduction

Neural signals such as electroencephalography (EEG), magnetoencephalography (MEG), electrocorticography (ECoG) and local field potential (LFP) provide critical insights into the physiological processes underlying key aspects of human cognition and neurodevelopment. These signals often reveal oscillations at different frequencies, which are represented as “bumps” in the power spectral density (PSD), and have been extensively studied as an objective measure for cognitive phenotyping (Klimesch, 1999; Schutter and Knyazev, 2012), biomarkers for age (Başar, 2013; Murty et al., 2020) as well as neurological disorders (Newson and Thiagarajan, 2018; Murty et al., 2021). In addition, the aperiodic background activity in the signals, often known as the ‘1/f component’ or ‘scale-free activity’, is characterized by the slope or exponent of the PSD (on a log-log scale) in a specified frequency band (He, 2014).

The PSD slope has garnered interest in recent years, as has been shown to be one of the key features of signal variability (Ribeiro and Castelo-Branco, 2022) and has been related to N900 lexical prediction (Dave et al., 2018), working memory (Donoghue et al., 2020b), grammar learning (Cross et al., 2022), sleep changes (Bódizs et al., 2021), anaesthesia (Kreuzer et al., 2020) and EEG fingerprinting (Demuru and Fraschini, 2020). The slope has been suggested to depend on multiple factors that include excitation-inhibition balance (Freeman and Zhai, 2009; Gao et al., 2017), dendritic response to an input (Voytek and Knight, 2015), tissue properties (Bédard et al., 2006a), temporal dynamics of the synaptic processes triggered by Poisson spiking (Milstein et al., 2009) and ionic diffusion processes across the extracellular membrane (Bédard and Destexhe, 2009). More recently, it has been linked to the “neural noise” hypothesis ((Voytek et al., 2015); more details in the Discussion section).

Change in aperiodic activity is suggested to be a plausible biomarker for some neurological and psychiatric diseases like schizophrenia (Molina et al., 2020), attention deficit hyperactivity disorder (ADHD; (Robertson et al., 2019; Ostlund et al., 2021)) and Fragile X Syndrome (Wilkinson and Nelson, 2021), although the aperiodic activity was unchanged in other diseases such as Alzheimer’s Disease (AD) (Benwell et al., 2020; Springer et al., 2022). Recently, reduction in the slope with increasing age has been observed across several studies under different task paradigms (Voytek et al., 2015; Dave et al., 2018; Tran et al., 2020) or even in resting state conditions (Thuwal et al., 2021; Hill et al., 2022; Merkin et al., 2023). However, slope analysis in these studies has been restricted to frequency ranges below 125 Hz, with most of them only up to 50 Hz. Indeed, very few reports have studied the PSD slopes in EEG at frequencies above 150 Hz (Dehghani et al., 2010; Moffett et al., 2017), but the effect of aging or mental disorder on the PSD slopes at these frequencies is unknown. However, the oscillations in high frequency range (300-500 Hz) also exhibit age dependant changes (Hume et al., 1982; Tanosaki et al., 1999), which prompted us to investigate the aperiodic activity in these frequency ranges.

In this study, we explored aperiodic activity in the resting state EEG activity up to 800 Hz in healthy elderly subjects aged 50-88 years. We analysed the slopes in both eyes open and eyes closed states. Eyes closed state enabled us to minimize potential electromyography (EMG) artifacts that could be present in eyes open state. We further examined the dependence of slopes on the reference schemes. Finally, we compared the slopes in subjects with mild cognitive impairment (MCI) and early AD with their age and gender matched healthy controls.

## Materials and Methods

The details of experimental setup and data collection have been explained in detail in previous studies (Murty et al., 2020, 2021; Kumar et al., 2022); here, we summarize them briefly.

### Dataset

We used the EEG dataset collected from 257 human subjects (109 females) aged 50-88 years (Murty et al., 2020, 2021) under Tata Longitudinal Study of Aging (TLSA) who were recruited from the urban communities of Bengaluru, Karnataka, India. They were clinically diagnosed by psychiatrists, psychologists and neurologists at National Institute of Mental Health and Neurosciences (NIMHANS), and M.S. Ramaiah Hospital, Bengaluru as cognitively healthy (N = 236), or with MCI (N = 15) or AD (N = 6), using a combination of tests such as Clinical Dementia Rating scale (CDR), Addenbrook’s Cognitive Examination-III (ACE-III), and Hindi Mental State Examination (HMSE). Diagnosis of all MCI/AD subjects was reviewed by a panel of four experts who reclassified two MCI subjects as healthy. Data from these two subjects were not taken for analyses. Further, 11 (10 healthy and 1 MCI) subjects were discarded due to noise (See Artifact Rejection subsection). One AD patient was further discarded as there was no age and gender matched control. We further discarded another 10 subjects (9 healthy and 1 MCI) for whom data was collected using 32 channels, leading to the usable 64-channel data of 233 subjects (217 healthy, 11 MCI and 5 AD). The eyes closed data (see the next subsection) was not collected from 16 subjects (13 healthy, 2 MCI and 1 AD) leading to a slightly smaller dataset for eyes closed condition.

Informed consent was obtained from the participants of the study and monetary compensation was provided. All the procedures were approved by The Institute Human Ethics Committees of Indian Institute of Science, NIMHANS, and M.S. Ramaiah Hospital, Bengaluru.

### Experimental Settings and Behavioural task

Briefly, EEG was recorded from 64-channel active electrodes (actiCap) using BrainAmp DC EEG acquisition system (Brain Products GMbH). The electrodes were placed according to the international 10–10 system, referenced online at FCz. Raw signals were filtered online between 0.016 Hz (first-order filter) and 1 kHz (fifth-order Butterworth filter), sampled at 2.5 kHz, and digitized at 16-bit resolution (0.1 μV/bit). The subjects were asked to sit in a dark room in front of a gamma corrected LCD monitor (BenQ XL2411; dimensions: 20.92 × 11.77 inches; resolution: 1280 × 720 pixels; refresh rate: 100 Hz) with their heads supported by a chin rest. It was placed at (mean ± SD) 58 ± 0.7 cm from the subjects (range: 54.9–61.0 cm) and subtended 52° × 30° of visual field for full screen gratings. Eye position was monitored using EyeLink 1000 (SR Research Ltd), sampled at 500 Hz.

In the beginning of the experiment, the resting state EEG for the eyes closed condition was collected for 1-2 minutes. Then, the subjects performed a passive visual fixation task, including the full screen grating stimuli. The task consisted of a single session that lasted for ~20 min, divided in 2–3 blocks with 3–5 min breaks in between, according to subjects’ comfort. Every trial started with the onset of a fixation spot (0.1°) shown at the centre of the screen, on which they were instructed to fixate. After an initial blank period of 1000 ms, 2–3 full screen grating stimuli were presented for 800 ms with an interstimulus interval of 700 ms using a customized software running on MAC OS. The stimuli were full contrast sinusoidal luminance achromatic gratings with either of the 3 spatial frequencies (1, 2, and 4 cycles per degree (cpd)) and 4 orientations (0°, 45°, 90°, and 135°). Our analyses were restricted to 500 ms of the interstimulus period before the onset of the stimulus, referred as the baseline period in the previous studies (Murty et al., 2020, 2021). We refer to it as “eyes open” condition.

### Artifact Rejection

For eyes open data, we used the artifact rejection framework as described in (Murty et al., 2020, 2021; Murty and Ray, 2022) and the steps are summarised here.

a. Eye-blinks or change in eye position outside a 5° fixation window during −0.5 s to 0.75 s from stimulus onset were noted as fixation breaks and removed offline. This led to a rejection of 14.6 ± 2.8% (mean ± SD) repeats.
b. All the electrodes with impedance >25 kΩ were rejected. Impedance of final set of electrodes was (mean ± SD) 5.48 ± 1.83 kΩ.
c. In the remaining electrodes, outliers were detected as repeats with deviation from the mean signal in (i) time or (ii) frequency domains by more than 6 standard deviations, and subsequently electrodes with more than 30% outliers were discarded.
d. Further, repeats that were deemed bad in the visual electrodes (P3, P1, P2, PO3, POz, PO4, O1, Oz and O2) or in more than 10% of the other electrodes were considered bad, eventually yielding a set of common bad repeats for each subject. Overall, this led us to reject (mean ± SD) 15.58 ± 5.48% repeats.
e. We computed slopes for the power spectrum between 56 Hz and 84 Hz range for each unipolar electrode and rejected electrodes whose slopes were less than 0. Overall, it led to a rejection of (19.45 ± 14.64%) electrodes.
f. We found that a small fraction of electrodes/stimulus repeats had either very small or very large signals that were not getting discarded using the pipeline above. Therefore, in addition to the artifact detection pipeline that was used in the previous studies, for each electrode, we computed the root mean square value (RMS) of the time series for all the remaining trials (after removing common bad repeats using the above criteria), and declared repeats having RMS values lower than 1.25 μV or higher than 35 μV as new outliers. We then rejected the electrodes that had more than 30% new outliers. Further, any new outlier if it belonged to the visual electrodes or was common in more than 10% of other electrodes was considered a bad repeat and appended to the existing list of bad repeats. This led to rejection of additional 1.24 ± 1.72 electrodes and 0.16 ± 1.05% repeats. This condition was added mainly to improve power spectral density (PSD) plots unlike the previous studies (Murty et al., 2020, 2021; Kumar et al., 2022) that dealt mainly with change in power. The main results remained similar even without applying this new criterion.
g. We further discarded the blocks which did not have at least a single good unipolar electrode in the left visual anterolateral (P3, P1, PO3, O1), right visual anterolateral (P2, P4, PO4, O2) and posteromedial (POz, Oz) electrode groups. We then pooled data across all good blocks for every subject separately for final analysis. Those subjects who did not have any analyzable blocks (10/237 healthy and 1/15 MCI) were discarded for further analysis.

The eyes closed EEG data was segmented into non-overlapping 2s epochs, resulting in a higher resolution of 0.5 Hz. Each segment was then treated as a “stimulus repeat” and subjected to the same artifact rejection pipeline as the eyes open data, with minor changes as described below. We only considered subjects who were deemed good for the eyes open condition. Then, we started with step (b) above, yielding the average impedance of 5.47 ± 1.77 kΩ. We then applied the RMS criteria in place of deviation from mean signal in time domain in (c). Since, we were applying RMS criteria at an initial step on the raw data containing the outliers, we had to increase the lower cut off to 2.5 μV and keep the upper cut off same. We repeated the remaining steps for artifact rejection done for eyes open from c (ii) - (e), i.e., detection of outliers from standard deviation in frequency domain to rejection of electrodes with slopes less than 0 as described above. It led to exclusion of (3.11% ± 4.87%) repeats and (18.86% ± 13.10%) electrodes. No additional subject was rejected when we applied the criteria in (g) for eyes closed data.

After discarding all the bad repeats, (293.42 ± 67.22) and (39.35 ± 7.75) repeats were available for eyes open and eyes closed conditions, respectively.

### EEG Data Analysis

Our primary emphasis was to characterize slope of the aperiodic activity in the PSD as a function of age within the elderly population (>49 years), for which we divided these subjects into two groups: 50–64 years (Mid) and >64 years (Old), as depicted in Table 1. We also compared the slopes in subjects with AD/MCI (termed “cases”) with their healthy, age and gender matched controls. As in our previous studies (Murty et al., 2021; Kumar and Ray, 2023), for each case, we averaged the relevant metrics for all age (± 1 year) and gender matched controls to yield a single control data point for each case, yielding 16 (13) pairs for eyes open (eyes closed) analyses (Table 1).

**Table 1:** Number of subjects in each group for the eyes open and eyes closed conditions. The numbers in parenthesis indicate the female subjects.

|  | Mid<br>(50-64 years) | Old<br>(>64 years) | MCI | AD |
| --- | --- | --- | --- | --- |
| Eyes open | 90 (50) | 127 (48) | 11 (2) | 5 (2) |
| Eyes closed | 82 (45) | 122 (48) | 9 (1) | 4 (2) |

All the data analyses were done using custom codes written in MATLAB (MathWorks. Inc; RRID:SCR_001622). Similar to the previous studies (Murty et al., 2020, 2021; Kumar et al., 2022) on this dataset, we chose [-500 0] ms as the eyes open period, yielding a resolution of 2 Hz. The analyses were performed using unipolar reference scheme, unless otherwise specified. Power spectrum was obtained using the Chronux Toolbox ((Bokil et al., 2010), RRID:SCR_005547) for individual trials and then averaged across the trials for each electrode.

#### Slope Analysis

The slope of the 1/f aperiodic component of the spectral distribution was computed using the Matlab wrapper for Fitting Oscillations and One Over f (FOOOF) toolbox (Donoghue et al., 2020b). In FOOOF, the power spectrum *P(f)* for frequency *f* is modelled as a combination of aperiodic *AP(f)* and oscillatory components and can be expressed as:

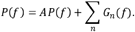

The *AP(f)* is given by

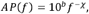

where *χ* is the exponent or ‘slope’ and *b* is the offset. Each oscillatory contribution *G_n_(f)* is modelled as a Gaussian peak:

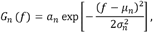

with *a_n_* as the peak height, *μ_n_* as the centre frequency and *σ_n_* as the width of each component. The settings chosen for FOOOF model parameters were: peak width limits: [4 8] for eyes open and [1 8] for eyes closed (different values were used to account for differences in frequency resolution in the two datasets); maximum number of peaks: 5; minimum peak height: 0.2; peak threshold: 2.0; and aperiodic mode: ‘fixed’.

In our data, peaks in the PSD were only evident in the alpha range (8-12 Hz) and line noise (50 Hz) and its harmonics. By taking a frequency range which did not include these oscillatory frequencies (or by not considering these frequencies for analysis), the slope can also be estimated by simply fitting a straight line to log(P(f)) versus log(f) plot, as done in a previous study (Shirhatti et al., 2016). We employed least square minimization using the program *fminsearch* in Matlab to obtain the slope, which gave similar results to the slopes estimated using FOOOF.

Since the slope does not remain constant throughout the full frequency range (Shirhatti et al., 2016), we fitted the PSD segments in frequency ranges from 4 to 1000 Hz in steps of 20 Hz with centre frequency between 40 Hz and 960 Hz, having the frequency width of 100 Hz. The PSD segments with the lowest and highest centre frequencies had slightly smaller frequency range width owing to frequency limit of 4-1000 Hz. We also fitted a single slope in the low frequency range (LFR) (64-140 Hz) and the high frequency range (HFR) (230-430 Hz) to verify that the trend in slope is not attributable to a particular frequency range width and length of PSD segments. Note that FOOOF has a limitation that if the frequency range to be fitted has peaks at its ends, the evaluated slope is inappropriate (Gerster et al., 2022). To overcome this limitation, we declared ±4 *Hz* around the peaks corresponding to the line noise at 50 Hz and its harmonics as “noise peaks”, and avoided frequency ranges for which these noise peaks lay near their end points. We also did not use slopes that were less than 0.01, which may be due to poor fitting.

#### Linear Regression

The slope was modelled to vary linearly with age. We used *fitlm* function in Matlab that generates the model parameters *β*1,*β*2 corresponding to *y* = *β*1 + *β*2 * *x*, R-squared (*R*^2^) and p-value using t-statistics.

#### Electrode Grouping

Scalp maps were generated using the topoplot function of EEGLAB toolbox ((Delorme and Makeig, 2004), RRID:SCR_007292) with standard *Acticap 64* unipolar montage of the channels. We divided the electrodes into 5 groups as occipital (O1, Oz, O2, PO3, PO4, PO7, PO8, PO9, PO10, POz), centro-parietal (CP1, CP2, CP3, CP4, CP5, CP6, CPz, P1, P2, P3, P4, P5, P6, P7, P8, Pz), fronto-central (FC1, FC2, FC3, FC4, FC5, FC6, C1, C2, C3, C4, C5, C6, Cz), frontal (Fp1, Fp2, F1, F2, F3, F4, F5, F6, F7, F8, Fz, AF3, AF4, AF7, AF8) and temporal (T7, T8, TP7, TP8, TP9, TP10, FT7, FT8, FT9, FT10). FCz and Fpz were used as the reference and the ground respectively. In our previous studies (Murty et al., 2020, 2021; Kumar et al., 2022), power analysis was done on electrodes for which strong gamma power was observed (P3, P1, P2, PO3, POz, PO4, O1, Oz and O2), which were termed as “high priority” electrodes. We performed slope analysis on this group as well.

### Statistical Analysis

The statistical analysis was done for slope comparison. Kruskal-Wallis (K-W) test, Wilcoxon rank sum (WRS) test were used to compare the medians across groups. The standard error of median (SEM) was computed after bootstrapping over 10,000 iterations.

To remove false alarms created by low p-values at certain frequencies, we applied cluster correction (Cohen, 2014). We used *bwconncomp* function of Image Processing toolbox in matlab to identify the frequency clusters having p-values less than 0.05 and 0.01. We called a cluster significant if at least three consecutive frequency bins were significant.

### Data and Code Availability

All spectral analyses were performed using Chronux toolbox (version 2.10), available at http://chronux.org. Slopes were obtained using matlab wrapper for FOOOF (https://github.com/fooof-tools/fooof_mat). Raw data will be made available to readers upon reasonable request and made publicly available at a later time.

## Results

We examined EEG data for 237 healthy adults aged between 50 and 88 years, grouped as Mid (50-64 years; N=90) and Old (>64 years; N=127). The subjects sat with their eyes closed for 1-2 minutes before performing a fixation task in which full-screen gratings were presented for 800 ms with an inter-stimulus interval of 700 ms. We computed the PSD of 500 ms segments of data during interstimulus period to get the “eyes open” condition, yielding a frequency resolution of 2 Hz. To overcome the plausible electromyography (EMG) artifacts in eyes open state, we also analysed the eyes closed data with segments of 2 s each, which yielded a higher resolution of 0.5 Hz. We computed the slopes using Matlab wrapper for Fitting Oscillations and One Over f (FOOOF) toolbox (Donoghue et al., 2020b), in steps of 20 Hz between 4 and 1000 Hz with centre frequencies from 40 Hz to 960 Hz by taking PSD segments of ± 50 Hz around each centre frequency. The first and the last PSD segments were slightly smaller owing to frequency limit of 4-1000 Hz. Slopes less than 0.01 were not considered for analysis. (See materials and methods). We have shown the PSDs and slopes up to 800 Hz to avoid the influence of filter roll off caused by a fifth order Butterworth filter at 1000 Hz.

### Slopes vary differently with age in low and high frequency regimes

Figure 1A shows the median PSDs and slopes with frequencies for the mid and old groups for the eyes open condition for the “high-priority” electrode group (see Methods for details). Like the slope variation in LFP data in monkeys (Shirhatti et al., 2016), for both age groups, slope was high initially due to the presence of oscillatory activities and decreased beyond ~60 Hz until ~140 Hz. It then increased again after a “knee” at ~175 Hz (see bottom plot in Fig. 1A). The PSDs flattened out for old subjects than mid aged subjects from ~50 Hz until ~140 Hz, resulting in lower slopes for the former in the low frequency range (LFR) 64-140 Hz, as highlighted by yellow boxes. This is in consensus with the results reported previously (Voytek et al., 2015; Merkin et al., 2023). However, beyond ~200 Hz the trend reversed and the PSDs for old subjects became steeper, resulting in steeper slopes beyond ~200 Hz up to ~700 Hz. The high frequency range (HFR) at 230-430 Hz, marked in orange, highlights the prominent region of higher slopes for the old group.

**Figure 1:**
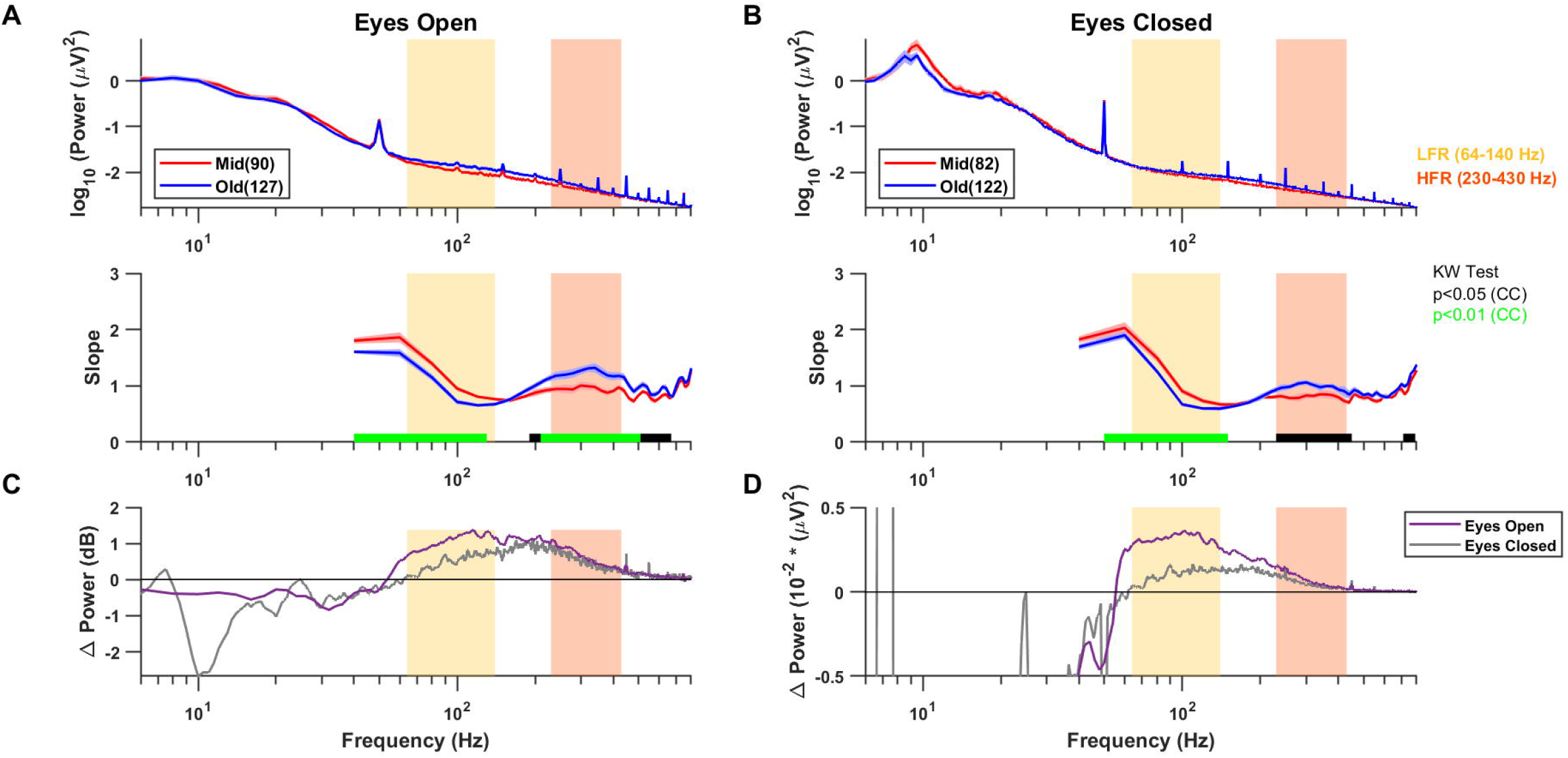
(A) PSDs (top) and slopes (bottom) for the two age groups in high priority electrodes in eyes open state. Solid traces represent the median and shaded region around them indicates ±SEM across subjects, computed after bootstrapping over 10,000 iterations. The numbers in legend in the top panel represent the subjects in the respective age groups. Coloured bars at the abscissa in the bottom panel represent significance of differences of slopes between mid and old (black: p<0.05 and green: p<0.01, K-W test, Cluster Corrected (CC)). Yellow and orange boxes represent the LFR (64-140 Hz) and the HFR (230-430 Hz) respectively. (B) Same as (A) in eyes closed condition. (C) The median change in power between mid and old in decibels (dB) for eyes open (purple) and eyes closed (grey) conditions. (D) Median change in absolute power between mid and old for these two conditions.

Since the frequency resolution for the eyes open condition was 2 Hz, and there could be some EMG activity related to maintenance of fixation, we also studied the slopes when eyes were closed for which longer segments of data were used to get a frequency resolution of 0.5 Hz (Fig. 1B). This condition revealed a more prominent alpha peak as compared to the eyes open condition, and also showed a well-known slowing of the alpha wave with aging (Scally et al., 2018; Merkin et al., 2023), with the centre frequency reducing from 9.72±0.11 in mid to 9.41 ±0.09 in old aged group (p=0.008, WRS test). Importantly, the flattening of slope in LFR and steepening in HFR with age was also observed in this dataset (Fig. 1B, bottom panel).

To better quantify the factors that could lead to the change in PSDs, we first compared the change in PSD between mid and old groups by subtracting the log PSD plots for the two conditions shown in Figures 1A and 1B (and multiplying by 10 to get units of decibels; Figure 1C). Since power is subtracted on a log scale, it is simply the log of the ratio of powers, and therefore, represents the scaling factor that must be applied on the PSD for the mid group to obtain the PSD for the old group. This log ratio was negative (scaling factor <1) below ~50 Hz, and subsequently became positive with a peak between 100-200 Hz. This ratio was smaller for the eyes closed condition in the LFR but comparable in the HFR.

We also considered an additive noise hypothesis in which noise was added to the signal obtained from the mid group to get the signal for the old group. If the noise at any frequency is independent of the signal, the power is additive, such that the power of the required noise can be obtained by simply subtracting the PSDs but on a linear (rather than log as done in Fig. 1C) scale (Fig. 1D). This also yielded a shallow “bump” between 50 Hz – 500 Hz peaking at around ~100 Hz (values below ~50 Hz are negative due to a reduction of alpha and beta power with aging). As in Figure 1C, noise power was lower in LFR for eyes closed than eyes open condition, potentially reflecting some contribution of EMG which is prominent between 40-200 Hz (Muthukumaraswamy, 2013; McManus et al., 2020). However, this “noise” was comparable in HFR range for the two conditions.

### Age variation of slope in LFR and HFR is observed in all electrodes

Next, we compared the slopes for different sensors, for which the electrodes were grouped in five categories as mentioned in Materials and Methods. Figure 2A shows the PSDs and slopes in all the electrode groups for eyes open condition. Interestingly, the PSD slopes varied considerably depending on electrode location over the entire frequency range. Figure 2C shows the fitted slopes in LFR and HFR. In particular, in HFR, the slopes increased radially from the central area.

**Figure 2.**
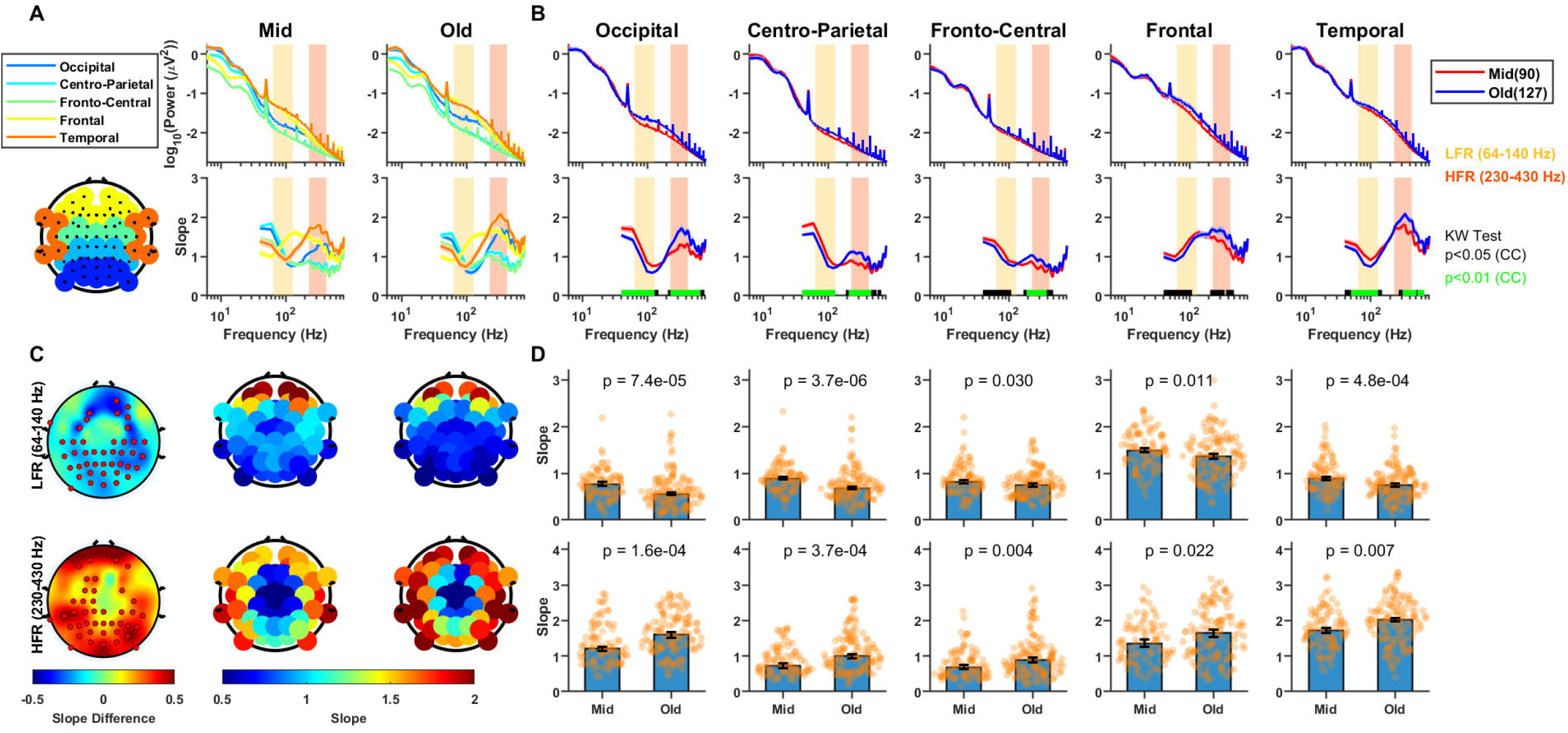
Variation of PSDs and slopes with frequency across the two age groups for different electrodes for the eyes open state. (A) PSDs and slopes across different electrode groups for mid and old age groups. The topoplot in the left bottom panel highlights the electrodes chosen for each electrode group. (B) PSDs and slopes between the age groups for each electrode group separately. Solid traces represent the median and shaded region around them indicates ±SEM across subjects, computed after bootstrapping over 10,000 iterations. The numbers in legend in the top panel represent the subjects in the respective age groups. Coloured bars at the abscissa in the bottom panel represent significance of differences of slopes between mid and old (black: p<0.05 and green: p<0.01, K-W test (CC)). Yellow and orange boxes represent the LFR (64-140 Hz) and the HFR (230-430 Hz) respectively in (A) and (B). (C) Topoplot of the difference in slopes between old and mid age groups (left column), computed in LFR (top row) and HFR (bottom row), and the raw slope values in the mid (central column) and old (right column) age groups. Electrodes with significant difference (p<0.05, WRS test) in slopes are highlighted in red in the left panel. (D) Bar plots showing median slopes for each electrode group for mid and old aged people for LFR and HFR, with error bars representing standard error of median (obtained using bootstrapping). The WRS test p-values are indicated at the top of the bar plots and the dots represent the slope for each subject.

In spite of the differences in absolute slope values, the difference in slope between old and mid aged groups showed consistent trends (Fig. 2B). The difference in slopes between old and mid groups in LFR (Fig. 2C; top plot) and HFR (Fig. 2C; bottom plot) were consistently negative and positive, respectively. The electrodes for which the differences were significant (p<0.05, Wilcoxon rank sum (WRS) test; red dots in Fig. 2C) were more concentrated in the posterior and occipital electrodes. Figure 2D shows the median value of the slopes in two age groups in LFR (top) and HFR (bottom), which were significantly different in all electrode groups (p-values are shown in the plots). The results were similar when the analysis was restricted to only males or females (data not shown). Similar analyses were performed for eyes closed state as shown in Figure 3. The results in eyes closed state were similar to the eyes open state, although the differences in slopes between the age groups (Fig. 3C) were less prominent and only significant in the occipital area for HFR (Figs. 3C and D).

**Figure 3.**
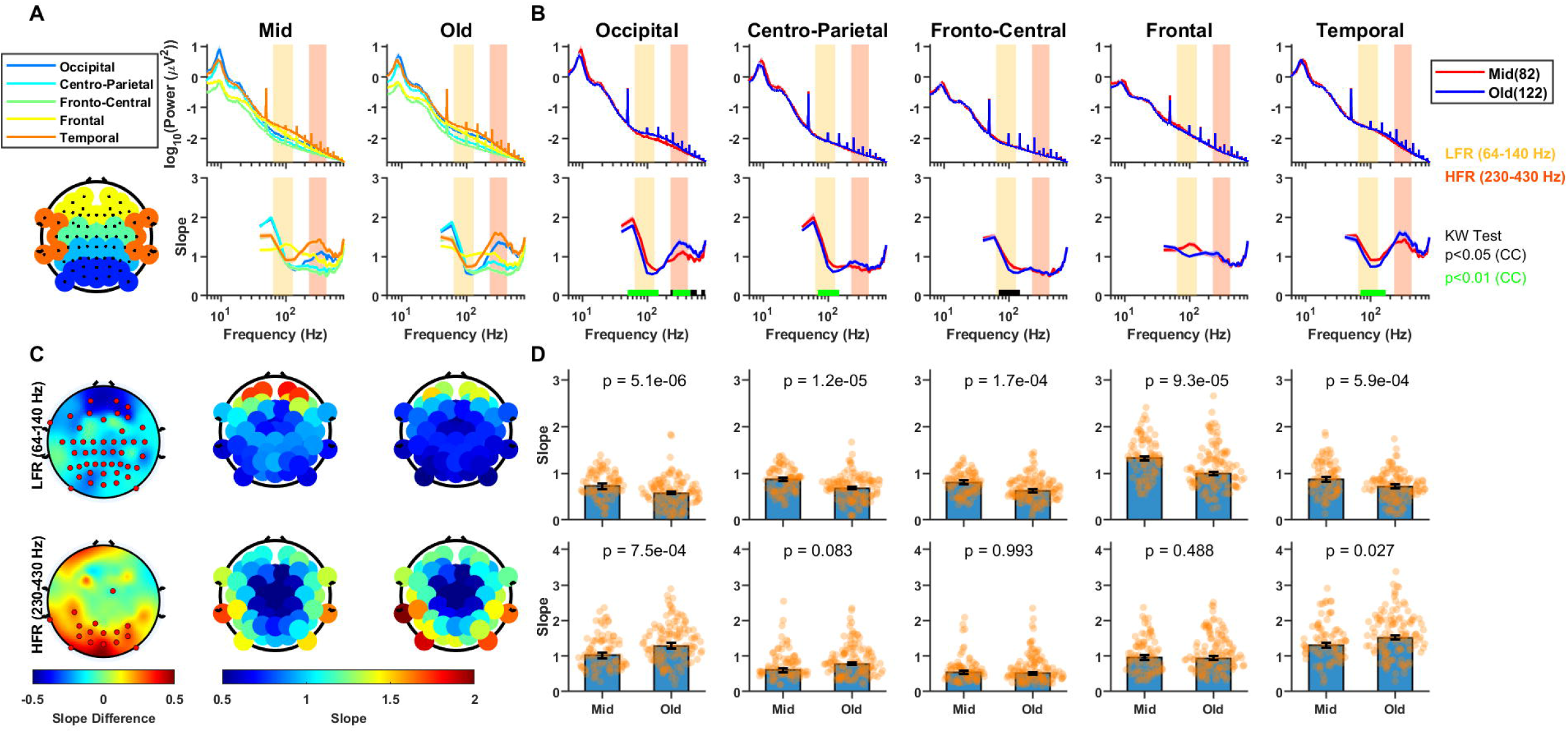
Variation of PSDs and slopes with frequency across the two age groups in various areas of brain for eyes closed state. Same as Figure 2 but for the eyes closed condition.

### Slope variation across electrodes is independent of the reference scheme

In HFR, the slopes were lower for the central electrodes and increased across frontal, temporal and occipital areas (Figs. 2C and 3C; bottom panels). In our recording setup, the reference and ground electrodes were near the centre, and therefore, the radial increase in slopes in HFR could be due to an increase with the distance of the electrode from the reference electrode. To test the dependence of HFR slope on the reference scheme, we re-referenced all signals using a bipolar referencing scheme in which each electrode was referenced with respect to a neighbouring electrode to yield a virtual bipolar electrode at the mid-point of the two electrodes, yielding 112 bipolar-referenced electrodes (for more details, see (Murty et al., 2020)). Figure 4 shows the same results as Figure 2C after computing the slopes for the bipolar signals. The results after bipolar referencing remained similar, thus, repudiating the variation across electrodes as an artifact, although the absolute slopes for bipolar reference scheme were slightly lower than unipolar reference scheme, as also shown previously (Shirhatti et al., 2016).

**Figure 4.**
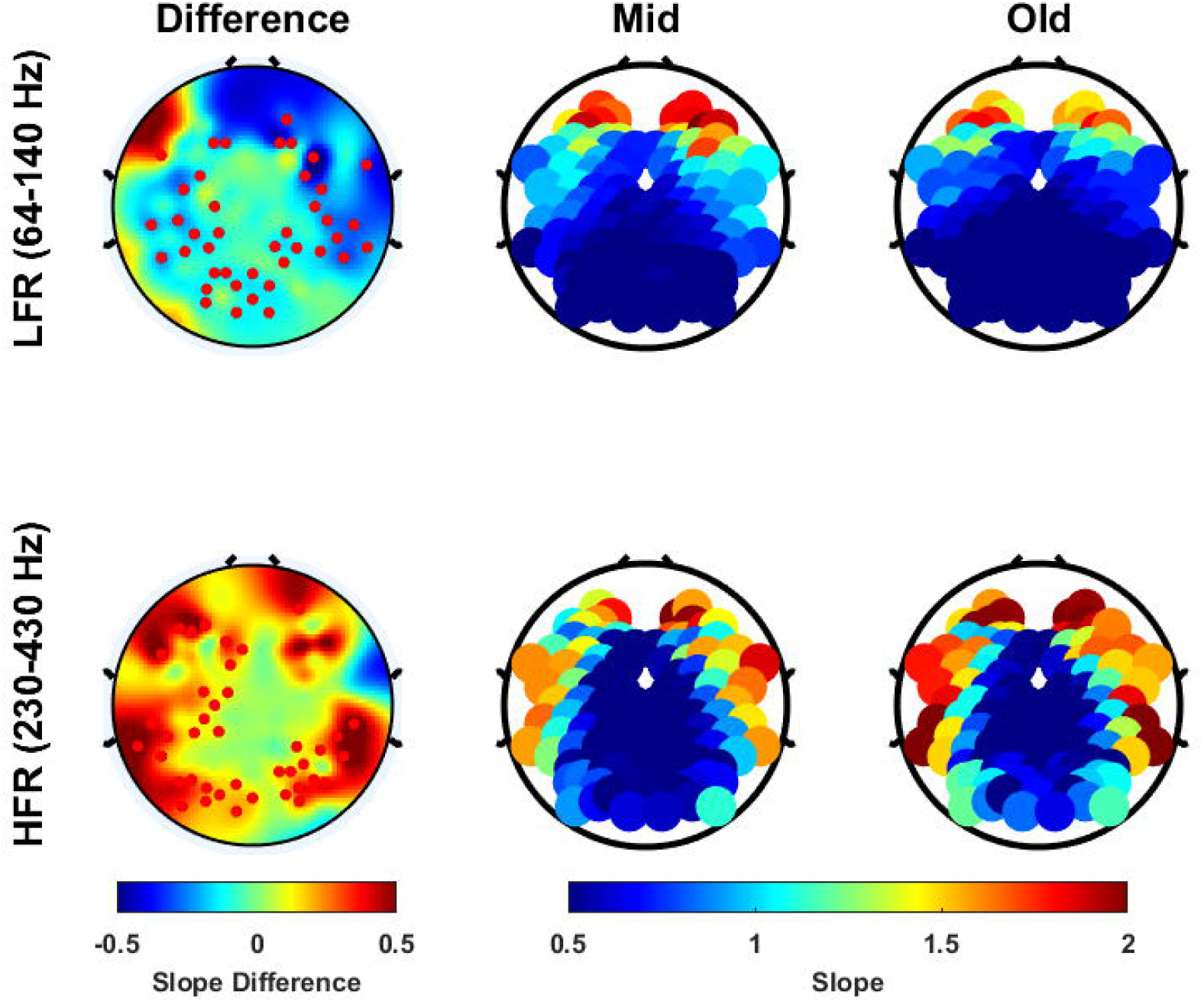
Same as Figure 2C but for bipolar reference scheme.

### Slopes do not vary between AD/MCI patients and their controls

We compared the PSDs and slopes in AD/MCI subjects with their age and gender matched healthy controls. As in our previous studies (Murty et al., 2021; Kumar and Ray, 2023), we averaged the PSDs/slopes across all controls for a given case subject to match the number of cases and controls. Figure 5 shows the results for the eyes closed condition (N=13). The PSDs revealed a significant reduction in beta power (15-35 Hz) in cases compared to controls in all the electrode groups as reported in other studies (Ranasinghe et al., 2022). For example, it reduced from 12.14±1.56 *μV*^2^ and 11.18±1.32*μV*^2^ in controls to 5.83±1.16 *μV*^2^ and 5.78 ±1.24 *μV*^2^ in cases in occipital and temporal electrodes (p=0.0048 and 0.0066, WRS test), respectively. This reduction was observed in the eyes open condition also (N=16), although it did not reach significance (data not shown). However, there was no significant change in slopes between controls and cases in either LFR or HFR in either eyes open and eyes closed conditions (eyes closed: Fig. 5B-D), consistent with previous studies which also did not find any changes in PSD slope with MCI/AD (Benwell et al., 2020; Springer et al., 2022).

**Figure 5.**
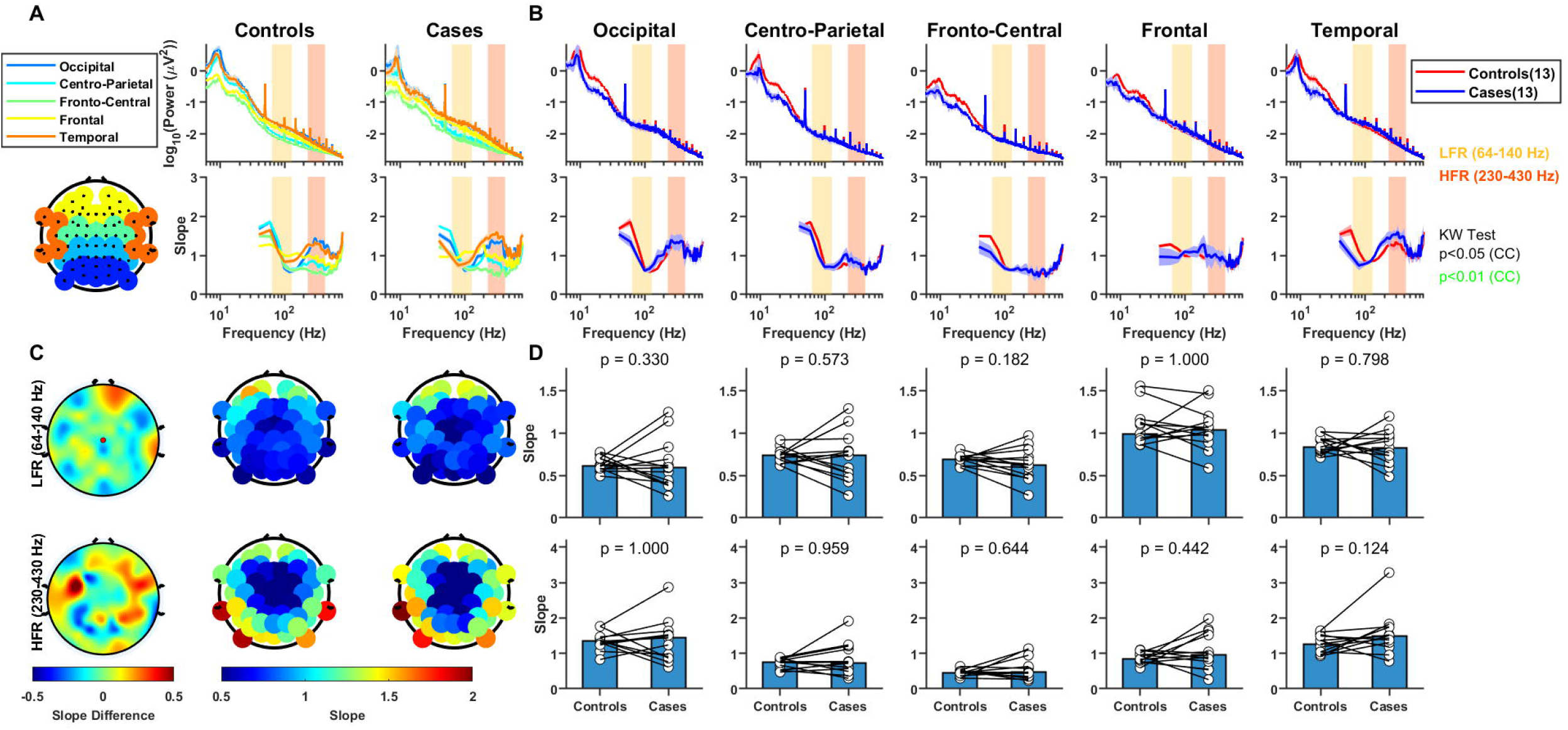
Comparison of PSDs and slopes with frequency between controls and cases (AD/MCI) for eyes closed state. (A)-(C) are similar to Figure 2A-C. Since there were several controls per case, median PSDs/slopes across all controls for each case subject was used to match the numbers of cases and controls. (D) The bar plots show the median slope for each electrode group in LFR and HFR across controls and cases. Individual case and control pair are shown as connected white dots. The WRS test p-values are indicated on the top of the bar plots.

### Variation in slope with age is confirmed using regression analysis

To confirm the variation in slope across subjects without grouping them in predefined groups, we regressed the slopes in the high priority electrode group with age (See Material and Methods for details) for the two frequency ranges and the two conditions, as shown in Figure 6. Consistent with previous results, we observed a negative relationship in LFR (slope=-0.010 for eyes open and −0.013 for eyes closed) and a positive relationship in HFR (slope=0.017 for eyes open and 0.016 for eyes closed). Interestingly, slope magnitude in HFR was higher than LFR, indicating a faster change of slope with age in HFR. Also, *R*^2^ was higher in HFR (0.53 eyes open, 0.46 eyes closed) than LFR (0.36 eyes open, 0.30 eyes closed). All the changes were significant (LFR: p=0.0007 eyes open, p=0.0001 eyes closed; HFR: p=0.0002 eyes open, p=0.0001 eyes closed). There was nearly equal distribution of AD/MCI cases, shown as pink dots, around the regression line indicating the indifference of their slopes with healthy subjects in both LFR and HFR for eyes open as well as eyes closed conditions.

**Figure 6.**
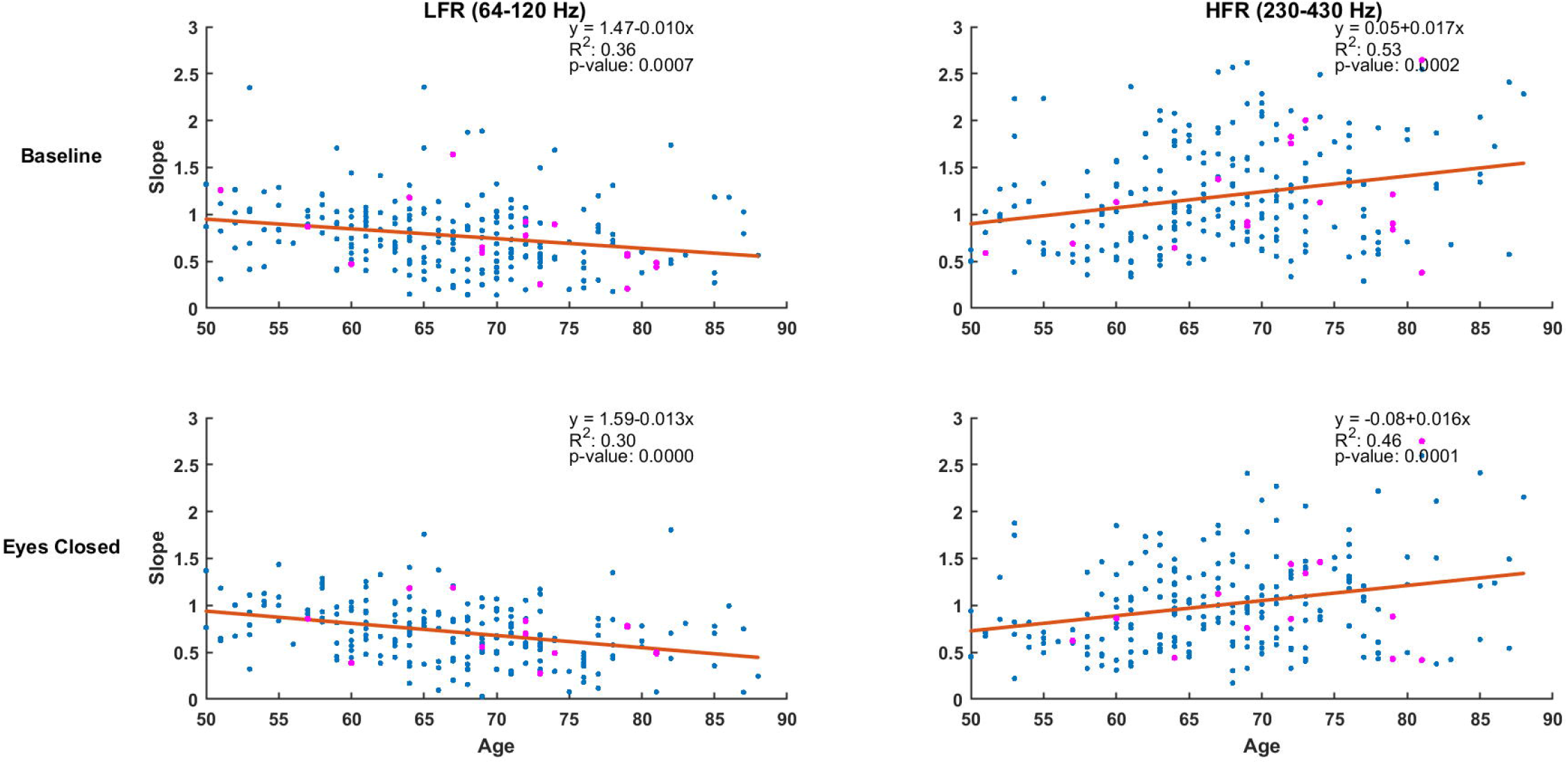
Scatter plots showing the slopes in high priority electrodes for all subjects in LFR and HFR for baseline and eyes closed conditions. Blue dots represent healthy subjects while MCI and AD subjects are indicated using pink dots. The regression line is in red and the corresponding parameters obtained using regression analysis are shown in top right side of each subplot.

## Discussion

We investigated age and MCI/AD related changes in the resting state aperiodic activity by studying the variation in the slope of the PSD over a broad frequency range (up to 800 Hz). In a task where subjects maintained fixation on the screen while visual stimuli were presented (Murty et al., 2020, 2021), we computed the PSDs during the inter-stimulus interval period. Consistent with previous studies, PSD slope flattened with age up to ~150 Hz. However, the slope showed a surprising increase with age at frequencies beyond 200 Hz. This age-related distinct behaviour of slopes at low and high frequency ranges was observed in all sensors, although, the slopes varied with sensor location. These results were confirmed using another dataset from the same subjects when they sat quietly with their eyes closed, for which PSDs could be obtained at higher frequency resolution. This ruled out the possibility of potential EMG artifacts related to open eyes or poor frequency resolution in the eyes open data affecting our results. The results remained similar when we used a bipolar reference scheme instead of the original unipolar reference scheme. Further, although we observed a reduction in beta power in subjects with MCI or AD compared to their age and gender matched healthy controls, we did not find any significant change in slopes.

### Previous studies on PSD slope changes with age

Voytek and colleagues first showed the reduction in the slope of the PSD in older (60-70 years) participants compared to younger (20-30 years) in a visual working memory task (Voytek et al., 2015), which has now been confirmed in several studies involving a task paradigm (Dave et al., 2018; Tran et al., 2020; Ribeiro and Castelo-Branco, 2022) as well as in resting state data (Hill et al., 2022; Merkin et al., 2023). However, all these studies dealt with frequencies less than 120 Hz, for example, 2-24 Hz (Voytek et al., 2015), 2-20 Hz (Tran et al., 2020), 1-40 Hz (Hill et al., 2022) and 2-40 Hz (Merkin et al., 2023). In our data, PSDs were largely overlapping up to ~50 Hz (Fig. 1), potentially because our analysis was limited to the elderly population between 50-90 years, as opposed to a wider gap in age groups in other studies (for example, 20-30 versus 60-70 in (Voytek et al., 2015) and 18-35 versus 50-86 years in (Merkin et al., 2023)). We therefore chose a slightly higher frequency range of 64-140 Hz (LFR) in which the PSD flattening could be easily observed in the PSDs, and 230-430 Hz (HFR) in which steepening of PSD was easily noticed.

### Neural mechanisms underlying PSD slope changes

Although precise generative mechanisms underlying aperiodic activity is unknown, a multitude of factors have been suggested to describe the slopes, that include tissue properties (Bédard et al., 2006a), self-organized criticality (SOC) (He et al., 2010; Chaudhuri et al., 2018), filtering properties of dendrites and extracellular medium (Bédard et al., 2006b; Logothetis et al., 2007; Bédard and Destexhe, 2009) and excitation-inhibition (E/I) balance (Gao et al., 2017; Naskar et al., 2021). In addition, the PSD slopes depend on simple factors such as the reference scheme used for recording the data (Shirhatti et al., 2016).

Recently, a “neural noise” hypothesis has been proposed to explain the reduction in the PSD slope, which is related with increased asynchronous background neuronal firing (Voytek et al., 2015) or an increase in E/I ratio (Gao et al., 2017). Specifically, Gao and colleagues developed a model in which Poisson distributed spikes arriving at excitatory and inhibitory neurons generated a change in excitatory and inhibitory conductance that were modelled as a difference-of-exponentials due to rise and decay time constants associated with AMPA and GABA_A_ receptors. Both these conductance profiles produced a low-pass effect, but slower kinetics of GABA_A_ caused a steeper roll-off with frequency compared to the AMPA conductance (see Fig. 1 of (Gao et al., 2017)). Consequently, reduction in inhibition (increase in E/I ratio), potentially due to dysfunctional inhibitory circuitry with aging, led to shallower slopes. This hypothesis has been supported by experiments in which the PSD slope was modulated by administration of drugs that led to either increased inhibition (e. g., propofol) or increased excitation (ketamine) (Gao et al., 2017; Lendner et al., 2020; Waschke et al., 2021). Our results could be consistent with this hypothesis, provided we modify the characteristics of this neural noise, which was band-pass as shown in Figure 1D.

Interestingly, surface EMG signals have energy between 40-200 Hz, peaking around ~100 Hz, which could explain at least part of the slope changes in the LFR (Muthukumaraswamy, 2013; McManus et al., 2020). On the other hand, this cannot explain the effect in steepening of the slope in HFR, since the noise was similar whether eyes were open or closed and occurred at a higher frequency range than EMG activity. Indeed, the noise need not be myogenic but instead generated by the brain. As the timescales of fundamental neural processes like synaptic neurotransmitter diffusion time and timing of spike propagation lie below 10 ms (Sabatini and Regehr, 1996; Shepherd, 2004), high frequency activity (>100 Hz) contains a wealth of information about neural processing and integration. Hence, the increase in slope in old group in HFR could be a reflection of age-related changes in these neural processes. In addition, it could be due to age-related alteration of physiological properties in brain tissue (Aalami et al., 2003; Lee and Kim, 2022), cortical thickness (Lillie et al., 2016) and skin conductance (Lim et al., 1996; Venables and Mitchell, 1996), which could determine the amplitude of this high-frequency signal measured from the scalp.

### Variation in PSDs slopes with frequency and electrode location

The PSD slopes in our study varied considerably depending on the frequency range (Fig. 1), similar to what we observed in LFP data in a previous study (see Fig. 1 of (Shirhatti et al., 2016)). We discuss some of these reasons below.

The initial high slope below 60 Hz as shown in Figures 1, 2, 3 and 5 may be due to the presence of the oscillatory activity that may have persisted even after accounting for the oscillatory power using FOOOF. The periodic activity in theta and alpha frequency range has been shown to be correlated with the aperiodic activity computed in the similar frequency range (Donoghue et al., 2020a). Also, researchers have stressed upon the need of removing the oscillations before computing the slopes indicating disruption of aperiodic activity in the low frequency ranges (Merkin et al., 2023). At lower frequencies, the slopes are also heavily dependent on the reference scheme (Shirhatti et al., 2016).

The dip in slope between 100-200 Hz and subsequent increase beyond 200 Hz reflects the presence of a “knee” in the PSD. This “knee” has been observed in previous studies as well, albeit at a different frequency. Miller and colleagues (Miller et al., 2009) found a knee in their ECoG PSD at ~75 Hz and attributed it to post-synaptic potential current of a particular timescale. In our data, the knee was present at ~175 Hz in all the electrode groups except the frontal group (where it was at ~100 Hz) and was prominent in occipital and temporal regions. This resulted in a timescale of ~5ms, computed as 1/2*πf_knee_*, *f_knee_* being the knee frequency. This timescale of ~5ms may be due to post-synaptic current, tissue low pass filter (Miller et al., 2009) or characteristic initial adaptation period of pyramidal neurons (Rauch et al., 2003).

The surprising result of the dependence of the PSD slope on electrode location at high frequencies, which has been observed previously (Dehghani et al., 2010), could be due to inhomogeneity of skull thickness or conductivity (Law, 1993; McCann et al., 2019). Future measurement of regional conductivity and its relation to PSD slope would be helpful to shed light on this issue.

### PSD slope changes with neurological disorders

Previous studies have shown a reduction in low-frequency oscillatory activity as well as stimulus-induced gamma activity in MCI/AD subjects (Benwell et al., 2020; Murty et al., 2021; Ranasinghe et al., 2022), which we also found in our dataset. Since neurological disorders are often associated with E/I imbalance (Lauterborn et al., 2021), and E/I ratio affects PSD slopes (Gao et al., 2017), we expected a change in PSD slope in MCI/AD subjects as well, but we found a null result. It could be due to a small sample size, although we found significant differences in power at both low-frequencies (Figure 5) and in the gamma band (Murty et al., 2021) with the same sample size. Further, this null result is consistent with previously documented MEG observations (Benwell et al., 2020; Springer et al., 2022). This suggests that the relationship between PSD slope and E/I balance could be complicated and dependent on other factors, as discussed above. Given that some other mental disorders have been shown to change the PSD slope (Molina et al., 2020), slope analysis could be used to distinguish between AD and other disorders. Future research related to how HFR slope varies with anaesthesia or other manipulations to change E/I balance will help elucidate its underlying mechanisms, and shed light on the neurobiology of aging in healthy and pathological brain.

## Funding disclosure and Acknowledgements

This work was supported by Tata Trusts Grant and Wellcome Trust/DBT India Alliance (Senior fellowship IA/S/18/2/504003) to SR) and Prime Minister Research Fellowship to SA.

## References

Aalami OO, Fang TD, Song HM, Nacamuli RP (2003) Physiological Features of Aging Persons. Archives of Surgery 138:1068–1076 Available at: https://doi.org/10.1001/archsurg.138.10.1068 [Accessed February 7, 2023].

Başar E (2013) A review of gamma oscillations in healthy subjects and in cognitive impairment. International Journal of Psychophysiology 90:99–117 Available at: https://www.sciencedirect.com/science/article/pii/S0167876013002134 [Accessed January 18, 2023].

Bédard C, Destexhe A (2009) Macroscopic Models of Local Field Potentials and the Apparent 1/f Noise in Brain Activity. Biophysical Journal 96:2589–2603 Available at: https://www.sciencedirect.com/science/article/pii/S0006349509004147 [Accessed January 19, 2023].

Bédard C, Kröger H, Destexhe A (2006a) Does the $l/f$ Frequency Scaling of Brain Signals Reflect Self-Organized Critical States? Phys Rev Lett 97:118102 Available at: https://link.aps.org/doi/10.1103/PhysRevLett.97.118102 [Accessed January 19, 2023].

Bédard C, Kröger H, Destexhe A (2006b) Model of low-pass filtering of local field potentials in brain tissue. Phys Rev E 73:051911 Available at: https://link.aps.org/doi/10.1103/PhysRevE.73.051911 [Accessed January 19, 2023].

Benwell CSY, Davila-Pérez P, Fried PJ, Jones RN, Travison TG, Santarnecchi E, Pascual-Leone A, Shafi MM (2020) EEG spectral power abnormalities and their relationship with cognitive dysfunction in patients with Alzheimer’s disease and type 2 diabetes. Neurobiol Aging 85:83–95.

Bódizs R, Szalárdy O, Horváth C, Ujma PP, Gombos F, Simor P, Pótári A, Zeising M, Steiger A, Dresler M (2021) A set of composite, non-redundant EEG measures of NREM sleep based on the power law scaling of the Fourier spectrum. Sci Rep 11:2041 Available at: https://www.nature.com/articles/s41598-021-81230-7 [Accessed January 19, 2023].

Bokil H, Andrews P, Kulkarni JE, Mehta S, Mitra PP (2010) Chronux: a platform for analyzing neural signals. J Neurosci Methods 192:146–151.

Chaudhuri R, He BJ, Wang X-J (2018) Random Recurrent Networks Near Criticality Capture the Broadband Power Distribution of Human ECoG Dynamics. Cerebral Cortex 28:3610–3622 Available at: https://academic.oup.com/cercor/article/28/10/3610/4508768 [Accessed January 22, 2023].

Cohen MX (2014) Analyzing neural time series data: theory and practice. Cambridge, Massachusetts: The MIT Press.

Cross ZR, Corcoran AW, Schlesewsky M, Kohler MJ, Bornkessel-Schlesewsky I (2022) Oscillatory and Aperiodic Neural Activity Jointly Predict Language Learning. Journal of Cognitive Neuroscience 34:1630–1649 Available at: https://doi.org/10.1162/jocn_a_01878 [Accessed January 19, 2023].

Dave S, Brothers TA, Swaab TY (2018) 1/f Neural Noise and Electrophysiological Indices of Contextual Prediction in Aging. Brain Res 1691:34–43 Available at: https://www.ncbi.nlm.nih.gov/pmc/articles/PMC5965691/ [Accessed January 19, 2023].

Dehghani N, Bédard C, Cash SS, Halgren E, Destexhe A (2010) Comparative power spectral analysis of simultaneous elecroencephalographic and magnetoencephalographic recordings in humans suggests non-resistive extracellular media. J Comput Neurosci 29:405–421 Available at: https://www.ncbi.nlm.nih.gov/pmc/articles/PMC2978899/ [Accessed February 7, 2023].

Delorme A, Makeig S (2004) EEGLAB: an open source toolbox for analysis of single-trial EEG dynamics including independent component analysis. J Neurosci Methods 134:9–21.

Demuru M, Fraschini M (2020) EEG fingerprinting: Subject-specific signature based on the aperiodic component of power spectrum. Computers in Biology and Medicine 120:103748 Available at: https://linkinghub.elsevier.com/retrieve/pii/S001048252030127X [Accessed January 19, 2023].

Donoghue T, Dominguez J, Voytek B (2020a) Electrophysiological Frequency Band Ratio Measures Conflate Periodic and Aperiodic Neural Activity. eNeuro 7 Available at: https://www.eneuro.org/content/7/6/ENEURO.0192-20.2020 [Accessed January 19, 2023].

Donoghue T, Haller M, Peterson EJ, Varma P, Sebastian P, Gao R, Noto T, Lara AH, Wallis JD, Knight RT, Shestyuk A, Voytek B (2020b) Parameterizing neural power spectra into periodic and aperiodic components. Nat Neurosci 23:1655–1665 Available at: https://www.nature.com/articles/s41593-020-00744-x [Accessed January 19, 2023].

Freeman WJ, Zhai J (2009) Simulated power spectral density (PSD) of background electrocorticogram (ECoG). Cogn Neurodyn 3:97–103.

Gao R, Peterson EJ, Voytek B (2017) Inferring synaptic excitation/inhibition balance from field potentials. NeuroImage 158:70–78 Available at: https://www.sciencedirect.com/science/article/pii/S1053811917305621 [Accessed January 19, 2023].

Gerster M, Waterstraat G, Litvak V, Lehnertz K, Schnitzler A, Florin E, Curio G, Nikulin V (2022) Separating Neural Oscillations from Aperiodic 1/f Activity: Challenges and Recommendations. Neuroinform 20:991–1012 Available at: https://link.springer.com/10.1007/s12021-022-09581-8 [Accessed January 20, 2023].

He BJ (2014) Scale-free brain activity: past, present, and future. Trends in Cognitive Sciences 18:480–487 Available at: https://www.sciencedirect.com/science/article/pii/S1364661314000850 [Accessed January 22, 2023].

He BJ, Zempel JM, Snyder AZ, Raichle ME (2010) The temporal structures and functional significance of scale-free brain activity. Neuron 66:353–369.

Hill AT, Clark GM, Bigelow FJ, Lum JAG, Enticott PG (2022) Periodic and aperiodic neural activity displays age-dependent changes across early-to-middle childhood. Developmental Cognitive Neuroscience 54:101076 Available at: https://linkinghub.elsevier.com/retrieve/pii/S1878929322000202 [Accessed January 19, 2023].

Hume AL, Cant BR, Shaw NA, Cowan JC (1982) Central somatosensory conduction time from 10 to 79 years. Electroencephalogr Clin Neurophysiol 54:49–54.

Klimesch W (1999) EEG alpha and theta oscillations reflect cognitive and memory performance: a review and analysis. Brain Research Reviews 29:169–195 Available at: https://www.sciencedirect.com/science/article/pii/S0165017398000563 [Accessed January 18, 2023].

Kreuzer M, Stern MA, Hight D, Berger S, Schneider G, Sleigh JW, García PS (2020) Spectral and Entropic Features Are Altered by Age in the Electroencephalogram in Patients under Sevoflurane Anesthesia. Anesthesiology 132:1003–1016 Available at: https://pubs.asahq.org/anesthesiology/article/132/5/1003/108947/Spectral-and-Entropic-Features-Are-Altered-by-Age [Accessed January 19, 2023].

Kumar WS, Manikandan K, Murty DVPS, Ramesh RG, Purokayastha S, Javali M, Rao NP, Ray S (2022) Stimulus-Induced Narrowband Gamma Oscillations are Test–Retest Reliable in Human EEG. Cereb Cortex Commun 3:tgab066 Available at: https://www.ncbi.nlm.nih.gov/pmc/articles/PMC8790174/ [Accessed November 29, 2022].

Kumar WS, Ray S (2023) Healthy aging and cognitive impairment alter EEG functional connectivity in distinct frequency bands.:2023.01.24.525301 Available at: https://www.biorxiv.org/content/10.1101/2023.01.24.525301vl [Accessed February 7, 2023].

Lauterborn JC, Scaduto P, Cox CD, Schulmann A, Lynch G, Gall CM, Keene CD, Limon A (2021) Increased excitatory to inhibitory synaptic ratio in parietal cortex samples from individuals with Alzheimer’s disease. Nat Commun 12:2603 Available at: https://www.nature.com/articles/s41467-021-22742-8 [Accessed February 10, 2023].

Law SK (1993) Thickness and resistivity variations over the upper surface of the human skull. Brain Topogr 6:99–109 Available at: https://doi.org/10.1007/BF01191074 [Accessed February 9, 2023].

Lee J, Kim H-J (2022) Normal Aging Induces Changes in the Brain and Neurodegeneration Progress: Review of the Structural, Biochemical, Metabolic, Cellular, and Molecular Changes. Frontiers in Aging Neuroscience 14 Available at: https://www.frontiersin.org/articles/10.3389/fnagi.2022.931536 [Accessed February 9, 2023].

Lendner JD, Helfrich RF, Mander BA, Romundstad L, Lin JJ, Walker MP, Larsson PG, Knight RT (2020) An electrophysiological marker of arousal level in humans Haegens S, Colgin LL, Piantoni G, eds. eLife 9:e55092 Available at: https://doi.org/10.7554/eLife.55092 [Accessed January 19, 2023].

Lillie EM, Urban JE, Lynch SK, Weaver AA, Stitzel JD (2016) Evaluation of Skull Cortical Thickness Changes With Age and Sex From Computed Tomography Scans. Journal of Bone and Mineral Research 31:299–307 Available at: https://onlinelibrary.wiley.com/doi/abs/10.1002/jbmr.2613 [Accessed February 8, 2023].

Lim CL, Barry RJ, Gordon E, Sawant A, Rennie C, Yiannikas C (1996) The relationship between quantified EEG and skin conductance level. International Journal of Psychophysiology 21:151–162 Available at: https://www.sciencedirect.com/science/article/pii/0167876095000496 [Accessed February 9, 2023].

Logothetis NK, Kayser C, Oeltermann A (2007) In vivo measurement of cortical impedance spectrum in monkeys: implications for signal propagation. Neuron 55:809–823.

McCann H, Pisano G, Beltrachini L (2019) Variation in Reported Human Head Tissue Electrical Conductivity Values. Brain Topogr 32:825–858 Available at: https://www.ncbi.nlm.nih.gov/pmc/articles/PMC6708046/ [Accessed February 9, 2023].

McManus L, De Vito G, Lowery MM (2020) Analysis and Biophysics of Surface EMG for Physiotherapists and Kinesiologists: Toward a Common Language With Rehabilitation Engineers. Front Neurol 11:576729 Available at: https://www.frontiersin.org/article/10.3389/fneur.2020.576729/full [Accessed February 8, 2023].

Merkin A, Sghirripa S, Graetz L, Smith AE, Hordacre B, Harris R, Pitcher J, Semmler J, Rogasch NC, Goldsworthy M (2023) Do age-related differences in aperiodic neural activity explain differences in resting EEG alpha? Neurobiology of Aging 121:78–87 Available at: https://www.sciencedirect.com/science/article/pii/S0197458022002019 [Accessed December 14, 2022].

Miller KJ, Sorensen LB, Ojemann JG, Nijs M den (2009) Power-Law Scaling in the Brain Surface Electric Potential. PLOS Computational Biology 5:e1000609 Available at: https://journals.plos.org/ploscompbiol/article?id=10.1371/journal.pcbi.1000609 [Accessed December 16, 2022].

Milstein J, Mormann F, Fried I, Koch C (2009) Neuronal Shot Noise and Brownian 1/f2 Behavior in the Local Field Potential. PLOS ONE 4:e4338 Available at: https://journals.plos.org/plosone/article?id=10.1371/journal.pone.0004338 [Accessed January 19, 2023].

Moffett SX, O’Malley SM, Man S, Hong D, Martin JV (2017) Dynamics of high frequency brain activity. Sci Rep 7:15758 Available at: https://www.ncbi.nlm.nih.gov/pmc/articles/PMC5693956/ [Accessed December 7, 2022].

Molina JL, Voytek B, Thomas ML, Joshi YB, Bhakta SG, Talledo JA, Swerdlow NR, Light GA (2020) Memantine Effects on Electroencephalographic Measures of Putative Excitatory/lnhibitory Balance in Schizophrenia. Biol Psychiatry Cogn Neurosci Neuroimaging 5:562–568.

Murty DV, Manikandan K, Kumar WS, Ramesh RG, Purokayastha S, Nagendra B, ML A, Balakrishnan A, Javali M, Rao NP, Ray S (2021) Stimulus-induced gamma rhythms are weaker in human elderly with mild cognitive impairment and Alzheimer’s disease Vinck M, Colgin LL, Bosman CA, eds. eLife 10:e61666 Available at: https://doi.org/10.7554/eLife.61666 [Accessed November 29, 2022].

Murty DVPS, Manikandan K, Kumar WS, Ramesh RG, Purokayastha S, Javali M, Rao NP, Ray S (2020) Gamma oscillations weaken with age in healthy elderly in human EEG. NeuroImage 215:116826 Available at: https://www.sciencedirect.com/science/article/pii/S105381192030313X [Accessed November 28, 2022].

Murty DVPS, Ray S (2022) Stimulus-induced Robust Narrow-band Gamma Oscillations in Human EEG Using Cartesian Gratings. Bio Protoc 12:e4379.

Muthukumaraswamy SD (2013) High-frequency brain activity and muscle artifacts in MEG/EEG: a review and recommendations. Front Hum Neurosci 7 Available at: http://journal.frontiersin.org/article/10.3389/fnhum.2013.00138/abstract [Accessed January 28, 2023].

Naskar A, Vattikonda A, Deco G, Roy D, Banerjee A (2021) Multiscale dynamic mean field (MDMF) model relates resting-state brain dynamics with local cortical excitatory–inhibitory neurotransmitter homeostasis. Network Neuroscience 5:757–782 Available at: https://doi.org/10.1162/netn_a_00197 [Accessed January 22, 2023].

Newson JJ, Thiagarajan TC (2018) EEG Frequency Bands in Psychiatric Disorders: A Review of Resting State Studies. Front Hum Neurosci 12:521.

Ostlund BD, Alperin BR, Drew T, Karalunas SL (2021) Behavioral and cognitive correlates of the aperiodic (1/f-like) exponent of the EEG power spectrum in adolescents with and without ADHD. Developmental Cognitive Neuroscience 48:100931 Available at: https://www.sciencedirect.com/science/article/pii/S1878929321000220 [Accessed January 19, 2023].

Ranasinghe KG, Verma P, Cai C, Xie X, Kudo K, Gao X, Lerner H, Mizuiri D, Strom A, Iaccarino L, La Joie R, Miller BL, Gorno-Tempini ML, Rankin KP, Jagust WJ, Vossel K, Rabinovici GD, Raj A, Nagarajan SS (2022) Altered excitatory and inhibitory neuronal subpopulation parameters are distinctly associated with tau and amyloid in Alzheimer’s disease Slutsky I, Chin J, Maestú F, eds. eLife 11:e77850 Available at: https://doi.org/10.7554/eLife.77850 [Accessed January 22, 2023].

Rauch A, La Camera G, Luscher H-R, Senn W, Fusi S (2003) Neocortical pyramidal cells respond as integrate-and-fire neurons to in vivo-like input currents. J Neurophysiol 90:1598–1612.

Ribeiro M, Castelo-Branco M (2022) Slow fluctuations in ongoing brain activity decrease in amplitude with ageing yet their impact on task-related evoked responses is dissociable from behavior O’Connell RG, Gold JI, O’Connell RG, Schölvinck M, eds. eLife 11:e75722 Available at: https://doi.org/10.7554/eLife.75722 [Accessed January 19, 2023].

Robertson MM, Furlong S, Voytek B, Donoghue T, Boettiger CA, Sheridan MA (2019) EEG power spectral slope differs by ADHD status and stimulant medication exposure in early childhood. J Neurophysiol 122:2427–2437.

Sabatini BL, Regehr WG (1996) Timing of neurotransmission at fast synapses in the mammalian brain. Nature 384:170–172 Available at: https://www.nature.com/articles/384170a0 [Accessed February 12, 2023].

Scally B, Burke MR, Bunce D, Delvenne J-F (2018) Resting-state EEG power and connectivity are associated with alpha peak frequency slowing in healthy aging. Neurobiol Aging 71:149–155.

Schutter DJLG, Knyazev GG (2012) Cross-frequency coupling of brain oscillations in studying motivation and emotion. Motiv Emot 36:46–54.

Shepherd GM (2004) The Synaptic Organization of the Brain. Oxford University Press. Available at: https://global.oup.com/academic/product/the-synaptic-organization-of-the-brain-9780195159561?cc=in&lang=en& [Accessed February 12, 2023].

Shirhatti V, Borthakur A, Ray S (2016) Effect of Reference Scheme on Power and Phase of the Local Field Potential. Neural Computation 28:882–913 Available at: https://direct.mit.edu/neco/article/28/5/882/8l59/Effect-of-Reference-Scheme-on-Power-and-Phase-of [Accessed January 19, 2023].

Springer SD, Wiesman AI, May PE, Schantell M, Johnson HJ, Willett MP, Castelblanco CA, Eastman JA, Christopher-Hayes NJ, Wolfson SL, Johnson CM, Murman DL, Wilson TW (2022) Altered visual entrainment in patients with Alzheimer’s disease: magnetoencephalography evidence. Brain Communications 4:fcac198 Available at: https://academic.oup.com/braincomms/article/doi/10.1093/braincomms/fcac198/6652979 [Accessed January 25, 2023].

Tanosaki M, Ozaki I, Shimamura H, Baba M, Matsunaga M (1999) Effects of aging on central conduction in somatosensory evoked potentials: evaluation of onset versus peak methods. Clin Neurophysiol 110:2094–2103.

Thuwal K, Banerjee A, Roy D (2021) Aperiodic and Periodic Components of Ongoing Oscillatory Brain Dynamics Link Distinct Functional Aspects of Cognition across Adult Lifespan. eNeuro 8:ENEURO.0224-21.2021 Available at: https://www.ncbi.nlm.nih.gov/pmc/articles/PMC8547598/ [Accessed December 7, 2022].

Tran TT, Rolle CE, Gazzaley A, Voytek B (2020) Linked sources of neural noise contribute to age-related cognitive decline. J Cogn Neurosci 32:1813–1822 Available at: https://www.ncbi.nlm.nih.gov/pmc/articles/PMC7474516/ [Accessed January 19, 2023].

Venables PH, Mitchell DA (1996) The effects of age, sex and time of testing on skin conductance activity. Biological Psychology 43:87–101 Available at: https://www.sciencedirect.com/science/article/pii/0301051196051836 [Accessed February 8, 2023].

Voytek B, Knight RT (2015) Dynamic Network Communication as a Unifying Neural Basis for Cognition, Development, Aging, and Disease. Biological Psychiatry 77:1089–1097 Available at: https://www.sciencedirect.com/science/article/pii/S0006322315003546 [Accessed January 19, 2023].

Voytek B, Kramer MA, Case J, Lepage KQ, Tempesta ZR, Knight RT, Gazzaley A (2015) Age-Related Changes in 1/f Neural Electrophysiological Noise. Journal of Neuroscience 35:13257–13265 Available at: https://www.jneurosci.org/lookup/doi/10.1523/JNEUROSCI.2332-14.2015 [Accessed January 19, 2023].

Waschke L, Donoghue T, Fiedler L, Smith S, Garrett DD, Voytek B, Obleser J (2021) Modality-specific tracking of attention and sensory statistics in the human electrophysiological spectral exponent Chait M, Shinn-Cunningham BG, Postle BR, Simon JZ, eds. eLife 10:e70068 Available at: https://doi.org/10.7554/eLife.70068 [Accessed January 19, 2023].

Wilkinson CL, Nelson CA (2021) Increased aperiodic gamma power in young boys with Fragile X Syndrome is associated with better language ability. Molecular Autism 12:17 Available at: https://doi.org/10.1186/s13229-021-00425-x [Accessed January 19, 2023].

